# Extensive and heterogeneous viral circular RNA production during Zika virus infection

**DOI:** 10.64898/2026.10.28.762464

**Authors:** Mirco Stradiotto, Alessandro Vezzi, Mariachiara Vardeu, Beatrice Mercorelli, Stefania Vogiatzis, Arianna Loregian, Luisa Barzon, Stefano Toppo, Enrico Lavezzo, Marta Trevisan

**Affiliations:** Department of Molecular Medicine, University of Padua, via Gabelli 63, 35121, Padova, Italy; Department of Biology, University of Padua, via Ugo Bassi 58b, 35121, Padova, Italy; Microbiology and Virology Unit, Padua University Hospital, via Giustiniani 1, 35128, Padova, Italy

**Keywords:** circRNAs, Zika virus, infection

## Abstract

Circular RNAs (circRNAs) are covalently closed RNA molecules with regulatory functions. Although virus-derived circRNAs (vcircRNAs) are well characterized in DNA viruses, their occurrence in RNA viruses remains poorly understood. Recent reports in coronaviruses and hepatitis C virus suggest that vcircRNA production extends beyond nuclear-replicating viruses. Here, we show that Zika virus (ZIKV), a vector-borne orthoflavivirus, produces vcircRNAs during infection of neural stem cells. Using a custom computational pipeline optimized for viral back-splice junction detection, we identified at 72 hours post infection a core set of 113 putative vcircRNAs shared across three biological replicates and distributed throughout the viral genome. Eight candidates spanning different genomic regions and abundance levels were experimentally validated by rolling-circle reverse transcription and Nanopore sequencing. These findings provide the first evidence of vcircRNA production in an orthoflavivirus, suggesting that circular RNA biogenesis may represent a previously underappreciated feature of RNA virus biology.

## Introduction

Circular RNAs (circRNAs) are a class of covalently closed RNAs whose functions in eukaryotic cells have only recently begun to emerge, including post-transcriptional regulation through microRNAs (miRNAs) sponging [1–3], protein decoy or scaffold functions [4,5], and serving as templates for peptide/protein translation [6]. Through these mechanisms, circRNAs can modulate key cellular processes, including cell differentiation and proliferation, cell cycle progression and apoptosis, senescence, autophagy, mitochondrial function, and protein homeostasis [7– 11]. CircRNAs are highly stable macromolecules, which present tissue- and cell-type specificity, and are particularly enriched in the brain [12]. Beyond eukaryotes, several DNA viruses express viral circRNAs (vcircRNAs), although functional roles have been identified for only a subset [13].

In contrast, RNA viruses would not be expected to generate vcircRNAs because circRNA biogenesis is typically linked to nuclear back-splicing of pre-mRNAs [1], a process absent from the predominantly cytoplasmic replication cycle of RNA viruses. Nevertheless, vcircRNAs have been identified in coronaviruses and in hepatitis C virus (HCV) [14,15], suggesting that circRNA production may represent a broader and previously underappreciated feature of RNA virus biology. Proposed functions for vcircRNAs produced by RNA viruses include the modulation of viral replication, host interactions, and pathogenesis [14,15].

Zika virus (ZIKV) is an arthropod-borne RNA virus belonging to the *Flaviviridae* family, genus O*rthoflavivirus*, that is associated with severe congenital abnormalities such as microcephaly and neurological complications [16,17]. In the developing brain, ZIKV preferentially infects neural stem cells (NSCs), impairing their survival and ultimately disrupting normal brain growth [18,19]. In 2024, the World Health Organization classified ZIKV as a high-priority pathogen with the potential to cause future Public Health Emergencies of International Concern because of its continued circulation across multiple countries [20]. Here, we report that ZIKV produces a large and heterogeneous population of vcircRNAs during infection of human NSCs, providing, to our knowledge, the first evidence of vcircRNA generation in an orthoflavivirus. The identification of these molecules required the development of a dedicated pipeline capable of detecting circRNAs derived from RNA viruses. A subset of these putative ZIKV vcircRNAs was experimentally validated using a novel experimental approach based on a combination of rolling circle reverse transcription and Nanopore sequencing. Overall, our findings extend vcircRNA biogenesis to orthoflaviviruses and further support the idea that circular RNA production may be a widespread feature of RNA viruses.

## Materials and methods

### Cell culture and viruses

Human neural stem cells (NSCs) were derived from H9 embryonic stem cells (hESCs, WAe009-A, WiCell Research Institute, Madison, WI) following a protocol previously described [21]. Briefly, H9 growing on Geltrex (Gibco) with mTeSR-1 (StemCell Technologies) medium were detached by Accutase (Sigma-Aldrich) treatment and seeded in 6-well Geltrex-coated plates in mTeSR-1 medium at a density of 2.5×10^5^ cells/well. The next day, Neurobasal medium supplemented with Neural induction supplement (both from Gibco) was added to the cells and was changed every other day from day 0 to day 7. Starting from day 7, NSCs were passaged with Accutase in NSCs-expansion medium (NEM) composed by 50% of Neurobasal medium, 50% of Advanced DMEM/F12 and 20% of Neural induction supplement (all from Gibco). Y-27632 (Selleck Chemicals) to a final concentration of 5 µM, was added at every passage. Vero cells and MRC-5 fibroblasts (ATCC) were cultured in Dulbecco modified essential medium (Gibco) supplemented with 10% fetal bovine serum (FBS, Gibco). All cells were maintained in an incubator at 37°C with 5% CO_2_ and 95% humidity.

Zika virus strain PRVABC59 (National Collection of Pathogenic Viruses, Public Health England, UK) was propagated and titrated, by standard plaque assay, on Vero cells.

### Infection experiments

NSCs were seeded in 6-well plates (5×10^5^) the day prior to the infection. Twenty-four hours later, cells were infected with ZIKV PRVABC59 at a multiplicity of infection (MOI) of 5 for 90 min, after which the virus was removed and replaced with regular NEM. Mock-infected NSCs were treated the same way except for the presence of the virus. At 24, 48 and 72 hours post-infection (hpi), mock and infected NSCs were harvested by Accutase treatment, washed with PBS (Gibco), pelleted and stored at −80 °C until RNA extraction. MRC-5 fibroblasts were seeded into 6 well plates (5×10^5^) 24 h before infection. The next day cells were infected with ZIKV at an MOI of 5 for 90 min. At 72 hpi, MRC-5 were harvested for subsequent analysis.

### RNA extraction

Ribonucleic acids were isolated from infected cells using the mirneasy tissue/cells Advance mini kit (Qiagen) following the manufacturer’s instructions. DNase (Qiagen) was applied in column during extraction. Total RNA was quantified with nanodrop and Qbit (both from Thermo Fisher Scientific). Total RNA quality was evaluated with the Agilent Tape station (Agilent).

### Sequencing

Illumina sequencing of RNA isolated from mock-infected and ZIKV-infected NSCs was outsourced to Novogene (Novogene Europe, 25 Science Park, Cambridge, CB4 0FW, UK), and performed using the company’s circRNA-seq protocol, “Eukaryotic circRNA (rRNA and linear RNA depletion)”.

### Bioinformatic identification and prioritization of viral circRNAs

Because commonly used circRNA detection tools rely on features of eukaryotic splice-site architecture that may not be applicable to RNA viruses, we developed a custom computational pipeline for the identification of virus-derived BSJs independently of canonical splice-site motifs. Following alignment filtering, non-colinear split reads were examined for back-splicing patterns, and candidate vcircRNAs were retained when supported by at least two BSJ-spanning reads.

To prioritize robust and reproducible events, candidates were clustered across biological replicates according to BSJ coordinates and subjected to a data-driven abundance filtering procedure based on normalized BSJ support, expressed as Counts Per Million Back-spliced Reads (CPMBR) (Supplementary Figure S1). Subsequent analyses focused on a core set of high-confidence vcircRNAs consistently detected across all three biological replicates at 72 hpi. Full details of BSJ detection, cross-replicate clustering, normalization and threshold selection are provided in the Supplementary Material.

### RT-qPCR analysis of ZIKV replication kinetics and selected ZIKV vcircRNAs

ZIKV replication kinetics and the expression of a subset of putative ZIKV vcircRNAs were analyzed by quantitative PCR (qPCR) using SYBR Green chemistry (Applied Biosystems). Total RNA was first reverse-transcribed using the SuperScript III Reverse Transcriptase kit (Invitrogen) according to the manufacturer’s instructions (Supplementary Material). qPCR was performed using SYBR Green master mix and primers targeting the ZIKV E gene or specific vcircRNA back-splice junctions (BSJs) (Supplementary Table S1), with a QuantStudio− 3 Real-Time PCR System (Applied Biosystems). ZIKV replication kinetics was expressed as 2^-ΔCt^, with Ct values normalized to the housekeeping gene *glyceraldehyde-3-phosphate dehydrogenase* (*GAPDH*). The relative abundance of vcircRNAs was expressed as log^10^(2−^ΔCt^), with Ct values normalized to the ZIKV E gene. Each experiment included at least three independent biological replicates (n ≥ 3).

### Rolling-circle RT and Nanopore validation of vcircRNAs

Selected vcircRNA candidates were experimentally validated using a long-read strategy combining rolling-circle reverse transcription (RRT), linear PCR amplification (L-PCR), and Nanopore sequencing (Fig. 2A). Total RNA was first polyadenylated with Poly(A) Polymerase (New England Biolabs, NEB) and treated with RNase R (Biosearch Technologies), to promote depletion of linear viral RNAs. Candidate-specific RRT was then performed using primers spanning the predicted BSJ (Supplementary Table S2), allowing reverse transcription to proceed iteratively around circular RNA templates and generate concatemeric cDNA products. Briefly, RRT products were initially amplified using a single candidate-specific primer located downstream of the BSJ. Because rolling-circle reverse transcription generates tandem repeats of the vcircRNA sequence, the resulting concatemeric cDNAs contain multiple copies of the L-PCR primer-binding site. Primer annealing at successive copies of this site can therefore generate amplification products encompassing one or multiple consecutive copies of the vcircRNA sequence. A second amplification step was subsequently performed using a candidate-specific reverse primer to generate double-stranded DNA (dsDNA) and amplification products were treated with Klenow fragment (NEB), to generate blunt-ended DNA molecules suitable for subsequent library preparation. The resulting dsDNA products were pooled together and sequenced on an Oxford Nanopore MinION platform (Oxford Nanopore Technologies), allowing the number of consecutive BSJ-spanning events to be resolved within individual long reads. Detailed reaction conditions and primer sequences are provided in the Supplementary Material.

### Bioinformatic analysis of Nanopore validation data

Nanopore reads generated from rolling-circle reverse transcription and linear amplification were analyzed against candidate-specific concatemeric references constructed by tandemly repeating the predicted vcircRNA sequence ten times and appending the corresponding linear-PCR primer sequence. Reads were aligned using minimap2 v2.30 [22] with the lr:hq preset and filtered to retain high-quality primary alignments. For each retained read, coverage across consecutive copies of the predicted circular sequence was evaluated relative to the RT-primer binding site, and the number of complete BSJ-spanning events was determined. Reads traversing the BSJ multiple times provided direct evidence of rolling-circle reverse transcription and were considered strong support for vcircRNA circularity. Detailed alignment filtering and BSJ-counting procedures are reported in the Supplementary Material.

## Data availability

All data and scripts required to reproduce the analyses presented in this study are publicly available on Zenodo under the reserved DOI [10.5281/zenodo.23390093].

## Results and Discussion

### ZIKV infection generates a reproducible core of viral circular RNA candidates

To investigate whether ZIKV generates viral circular RNAs (vcircRNAs) during infection, human neural stem cells (NSCs) were infected with ZIKV and collected at 24, 48 and 72 hours post-infection (hpi). Following enrichment for circular RNAs, samples were subjected to paired-end RNA sequencing (Fig. 1A and Supplementary Table S3) and analyzed using a custom back splice junction (BSJ)-detection pipeline, enabling the identification of putative viral BSJs in ZIKV-infected samples (Supplementary Table S4). Viral RNA levels progressively increased during infection, with a 94.5-fold increase from 24 to 72 hpi (Fig. 1B). Notably, the proportion of viral reads spanning putative BSJs also increased over time, from approximately 1% at 24 hpi to 3% at 72 hpi (Fig. 1C). Because BSJ-spanning reads were normalized to total viral reads, this trend cannot be solely attributed to the overall increase in viral RNA abundance but indicates a progressive enrichment of BSJ-containing RNAs within the viral RNA population during infection.

**Figure 1.**
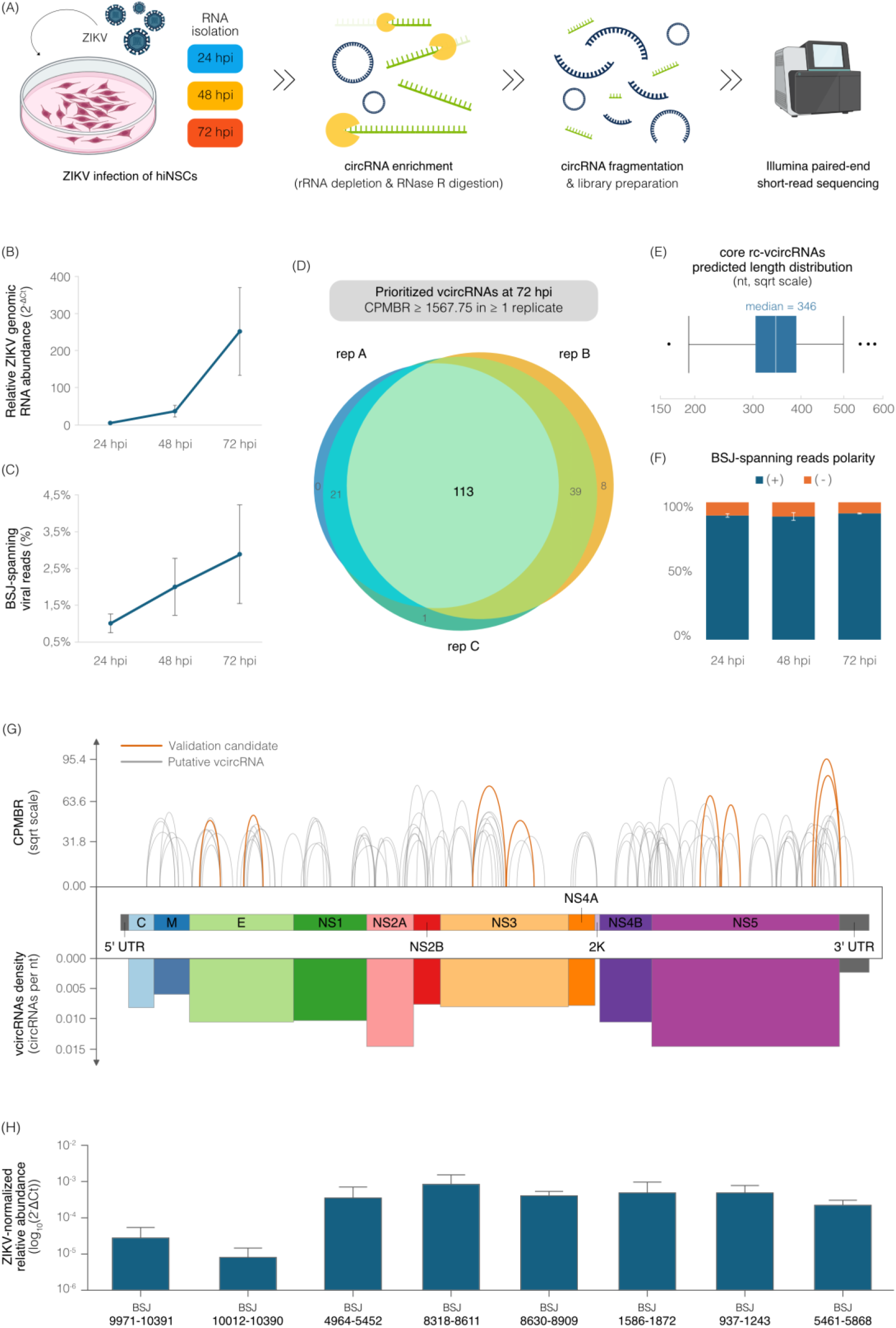
Identification, characterization, and validation of Zika virus (ZIKV)-derived circular RNAs in infected neural stem cells (NSCs). (A) Experimental workflow for virus-derived circRNA (vcircRNA) detection in ZIKV-infected NSCs at 24, 48 and 72 hpi. Created with BioRender. (B) Viral RNA abundance at 24, 48, and 72 hpi, expressed as relative RNA levels normalized to *glyceraldehyde-3-phosphate dehydrogenase* (*GAPDH*) housekeeping gene (mean ± SD, n=3 biological replicates). (C) Percentage of ZIKV-mapped reads supporting back-splice junctions (BSJs) relative to the total number of reads mapped to the ZIKV genome at each time point (mean percentage ± SD, n=3 biological replicates) (D) Distribution of prioritized vcircRNAs across three biological replicates at 72 hpi. Candidates were retained if their expression reached a threshold of Count Per Million Backspliced Reads (CPMBR) ≥ 1567.75 in at least one replicate, while detection in individual replicates was defined as CPMBR > 0. The 113 vcircRNAs shared across all three replicates were designated as core replicate-consistent vcircRNAs (rc-vcircRNAs). (E) Predicted length distribution of core rc-vcircRNAs. (F) Proportion of BSJ-spanning reads with positive (+) and negative (-) polarity at each time point. (G) Genomic distribution of core rc-vcircRNAs along the ZIKV genome, with candidate vcircRNAs selected for experimental validation highlighted in orange. The lower panel shows vcircRNA density across the viral genome, together with the annotated viral genomic regions. (H) Candidate core rc-vcircRNAs validation by quantitative reverse-transcription PCR (RT-qPCR). Relative abundance is expressed as log^10^(2−^ΔCt^), with Ct values normalized to ZIKV genomic RNA quantified using primers targeting the E gene. Data show the mean ± SD of n = 3 independent experiments.

**Figure 2.**
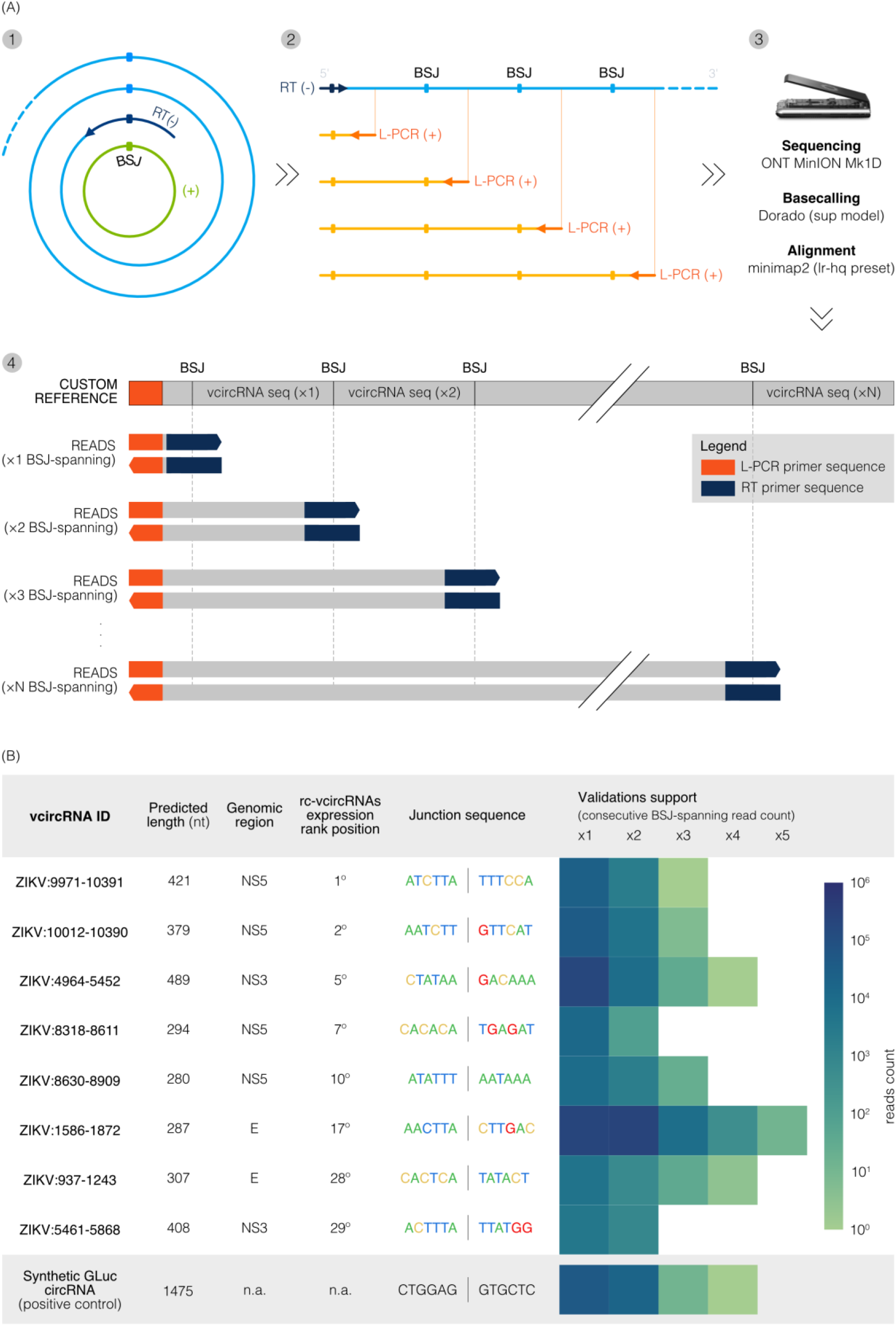
Candidate core replicate-consistent vcircRNA (rc-vcircRNA) validation through rolling-circle reverse transcription (RRT), linear PCR (L-PCR) and Nanopore sequencing. **(A)** Schematic overview of the validation workflow. **1:** An RT primer spanning the BSJ enables RRT of rc-vcircRNAs. **2:** Concatemeric amplicons containing multiple consecutive BSJ repeats were generated from rolling-circle RT products using a single L-PCR primer downstream of the BSJ. **3:** Long-read sequencing was performed using the Oxford Nanopore MinION Mk1D platform, followed by basecalling (Dorado, sup model) and alignment to a custom reference using minimap2 (lr:hq preset). **4:** The custom reference was generated by concatenating 10 copies of the rc-vcircRNA sequence defined by the predicted coordinates. An L-PCR primer sequence was added at the beginning of the reference to match the expected read structure. Reads containing at least two consecutive BSJ crossings were considered strong evidence of rc-vcircRNA validation. **(B)** Summary of the eight rc-vcircRNA candidates selected for experimental validation, including their genomic coordinates, predicted length, genomic region, expression-based ranking among core rc-vcircRNAs, and BSJ sequences. The heatmap shows read counts stratified by the number of consecutive BSJ crossings (one to five), with colors representing counts on a logarithmic scale. Validation results for the synthetic *Gaussia* luciferase (GLuc) circRNA, used as a positive control, are also reported.

One possible explanation for this progressive enrichment may be the greater stability of circRNAs: their covalently closed structure, lacking free 5′ and 3′ ends, makes them more resistant to exonucleolytic degradation and may allow them to persist longer than linear RNAs [23–25]. Thus, the increasing relative abundance of ZIKV BSJ-containing RNAs over time could reflect, at least in part, preferential accumulation of circular molecules during infection rather than an increased rate of circular RNA biogenesis. Direct measurements of vcircRNA turnover will be required to distinguish between these possibilities.

Initial BSJ detection revealed a highly heterogeneous landscape dominated by events supported by few sequencing reads. To focus subsequent analyses on the most robust candidates, we selected the 72 hpi time point, which exhibited the highest viral RNA abundance, and applied within-replicate denoising followed by a data-driven abundance and reproducibility filtering procedure designed to preferentially retain consistently supported BSJs. This approach identified 113 high-confidence vcircRNA candidates shared across all three biological replicates, together with smaller sets supported by two replicates (Fig. 1D). We refer to this reproducible set of candidates shared across all replicates as the core of replicate-consistent vcircRNAs (core rc-vcircRNAs).

Core rc-vcircRNAs displayed substantial size heterogeneity, ranging from approximately 160 to 580 nt, with a median predicted length of 346 (Fig. 1E). This size distribution substantially overlaps with the range reported for HCV-derived circRNAs, (approximately 130 nt to >2,000 nt), most of which are a few hundred nucleotides long (200-300 nt) [14].

BSJ-spanning reads showed a striking and consistent strand bias throughout the infection, with approximately 90% of them originating from the positive-sense strand (Fig. 1F). A similar positive-strand predominance has been reported for HCV vcircRNAs, for which ∼98% of junction-spanning reads derive from the positive-sense genome [14]. The concordant strand preference observed in two distantly related members of the *Flaviviridae* family is intriguing, but whether it might reflect the greater abundance and accessibility of positive-sense genomic RNA or a specific feature of the circularization process remains to be established.

Core rc-vcircRNA candidates mapped throughout the ZIKV genome, encompassing both structural and non-structural coding regions and untranslated regions, displaying substantial differences in normalized BSJ abundance (Fig. 1G). Although several relatively abundant candidates were observed, no single genomic region accounted for the overall population. A similarly broad distribution of circularization events has been reported for HCV and coronaviruses, where BSJs occur throughout the viral genome while showing local clustering at preferential breakpoint regions [14,15,26]. Notably, closely related ZIKV BSJs frequently displayed start and end coordinates differing by only a few nucleotides, indicating substantial heterogeneity in breakpoint selection. A remarkably similar pattern has been experimentally observed for SARS-CoV-2 vcircRNAs, where individual vcircRNA species displayed multiple closely spaced BSJ variants rather than uniquely defined junction coordinates [15,26]. This shared breakpoint flexibility contrasts with the precise splice-site selection characteristic of canonical spliceosome-dependent back-splicing and may therefore reflect a distinct, potentially error-prone mechanism of circular RNA formation in cytoplasmic RNA viruses.

Overall, these data identify a reproducible population of ZIKV-derived BSJ-containing RNAs that progressively becomes enriched during infection, originates predominantly from the positive-sense genome and encompasses diverse regions of the viral RNA. Their broad distribution and heterogeneous junction architecture raise the question of whether these BSJs originate from *bona fide* circular RNA molecules rather than from other non-colinear viral RNA species.

### Experimental validation confirms ZIKV BSJs and provides direct evidence of RNA circularity

We next sought to independently confirm selected BSJs and subsequently obtain direct structural evidence of RNA circularity. Eight core rc-vcircRNA candidates were selected from the top 30 most abundant candidates, spanning different abundance ranks, predicted sizes and genomic regions (E, NS3, and NS5), thereby providing a representative panel of the high-confidence vcircRNA population. Divergent primers flanking the predicted BSJs were used to quantify the selected candidates by RT-qPCR following reverse transcription with random hexamers. All eight candidates were detected in ZIKV-infected cells, with ZIKV-normalized relative abundances spanning approximately two orders of magnitude (Fig. 1H). Thus, targeted RT-qPCR independently confirmed the detection of all eight vcircRNA-seq-derived candidates, which were also detected in ZIKV-infected MRC-5 fibroblast cells (Fig. S2), indicating that their expression is not restricted to NSCs. This finding suggests that ZIKV vcircRNA expression may occur across different cell types.

BSJ detection using divergent primers does not, by itself, establish the circular nature of the corresponding RNA molecule. To provide more direct molecular evidence of RNA circularity, we developed a long-read validation strategy combining rolling-circle reverse transcription (RRT), linear amplification, and Nanopore sequencing (Fig. 2A). Candidate-specific primers spanning the predicted BSJ were used to initiate reverse transcription. On a circular template, continued reverse transcription around the molecule generates concatemeric cDNAs containing tandem copies of the vcircRNA sequence and, consequently, repeated copies of the same BSJ. These concatemeric products were amplified, converted to double-stranded DNA and subjected to Nanopore sequencing, enabling repeated BSJ traversals to be resolved within individual long reads.

Application of this approach to the same eight core rc-vcircRNAs generated long reads repeatedly traversing the predicted BSJ for every candidate (Fig. 2B). A synthetic circular RNA encoding the *Gaussia* luciferase (GLuc) was also included as a positive control to assess the ability of the workflow to generate and detect concatemeric products from a circular RNA template. Remarkably, reads traversing the predicted BSJ at least two consecutive times were recovered for all eight vcircRNAs, with individual molecules containing up to five consecutive BSJ-spanning events. The recovery of individual long reads containing repeated copies of the same non-colinear junction provides strong structural evidence supporting the circular nature of the corresponding ZIKV-derived RNA molecules.

Interestingly, the same approach also provided structural support for four additional vcircRNA candidates that fell outside the stringent criteria used to define the core set (Supplementary Fig. S3). Although these candidates were not considered in the primary analyses, their independent validation suggests that the conservative filtering strategy may exclude genuine vcircRNAs with lower or more variable support across short-read libraries.

Together, circRNA-enriched short-read sequencing, targeted RT-qPCR, and rolling-circle long-read validation provide convergent evidence for ZIKV-derived circular RNAs. Structural validation across different abundance ranks and genomic regions, together with additional validation of non-core candidates, indicates that ZIKV vcircRNA landscape is not restricted to a small number of highly abundant or recurrent species.

Collectively, our findings reveal a diverse population of ZIKV-derived circular RNAs and extend vcircRNA formation to the *Orthoflavivirus* genus. Their heterogeneous size and breakpoint architecture, broad genomic distribution, and progressive enrichment during infection reveal an additional layer of complexity within the ZIKV RNA population. Together with previous observations in HCV and coronaviruses, these findings suggest that circular RNA formation may represent a broader feature of cytoplasmic RNA virus biology. Whether these molecules arise as by-products of viral RNA replication or exert specific biological functions remains to be determined. Finally, to support future investigations, we developed a versatile framework combining circRNA-seq with Nanopore sequencing for circRNA discovery and single-molecule structural validation across diverse biological contexts.

## Acknowledgments

This study was supported by University of Padua, Italy (PRID 2022) to M.T., by the DURABLE project, co-funded by the European Union, under the EU4Health Programme (EU4H), project no. 101102733, and by the REFINE project, co-funded by the European Union, under the Horizon Europe Programme, project n. 101191791.

