## Supplementary material for "Extensive and heterogeneous viral circular RNA production during Zika virus infection"

**Supplementary methods**

**Supplementary Figures S1-S3**

**Supplementary Tables S1-S4**

### Supplementary Methods

#### Computational detection of viral circular RNAs

*Alignment filtering and identification of split reads*

A custom computational pipeline was developed to identify virus-derived back-splice junctions (BSJs) independently of canonical splice-site motifs. The approach relies on non-colinear alignment patterns generated by reads spanning putative circular RNA junctions.

Aligned reads in SAM format were initially filtered to remove unmapped reads and reads flagged as PCR or optical duplicates. Then, to identify unambiguous alignments, read pairs were filtered to retain a clear best alignment configuration for each mate. Alignments were grouped by query name, and mates were evaluated independently. Candidate configurations consisted of either a single non-supplementary alignment or a non-supplementary alignment paired with a supplementary alignment, provided that both had the same secondary-alignment flag status. Candidates were ranked by summed alignment scores (AS), penalized for excessive query overlap, and filtered by minimum query coverage (95%). The best configuration was accepted if it was the only eligible candidate or exceeded the second-best score by the required margin (min = 5); mates with a single alignment were accepted directly. Read pairs were retained only if both mates passed, and only the selected alignments were reported. Supplementary alignments have been considered since they represent additional alignment segments of split reads and are therefore required to reconstruct non-colinear mapping patterns. Following this initial filtering, only reads associated with at least one supplementary alignment were considered for BSJ detection.

Primary and supplementary alignment segments belonging to the same read were grouped and ordered according to their query coordinates. Consecutive segments were then examined for consistency between their order along the read and their genomic coordinates. A non-colinear configuration compatible with back-splicing was identified when a downstream segment along the read mapped upstream of the preceding segment on the viral reference genome. A tolerance of up to 5 nt was allowed when evaluating coordinate inversion to account for local alignment uncertainty and short sequence repetitions around candidate junctions.

*Paired-end support and within-replicate denoising*

BSJ detection was performed independently for each read of a paired-end fragment. A read pair was considered to support a candidate vcircRNA when one read displayed a non-colinear split alignment compatible with a BSJ and its mate mapped within the genomic interval delimited by the inferred circular RNA boundaries.

Mate support required both the start and end coordinates of the mate alignment to fall within the predicted vcircRNA interval, allowing 5 nt tolerance beyond either boundary. When both reads of a pair independently spanned a putative BSJ, junction coordinates inferred from the two reads were compared and junctions differing by no more than 2 nt were considered concordant.

Candidate BSJs were initially identified independently within each biological replicate. To reduce the contribution of isolated events with minimal sequencing support, candidates supported by a single BSJ-spanning read within a replicate were removed before cross-replicate comparison.

#### Cross-replicate clustering, normalization and definition of the high-confidence vcircRNA core

*Cross-replicate clustering*

Candidate BSJs passing within-replicate denoising were compared across biological replicates according to their junction coordinates. To account for minor variability in alignment boundaries, BSJs were considered equivalent when their start and end coordinates matched within a tolerance of ± 5 nt. Equivalent junctions were grouped into cross-replicate clusters, allowing evidence for closely matching BSJs to be integrated across biological replicates.

For each cluster, BSJ support was quantified independently in each replicate and normalized as Counts Per Million Back-spliced Reads (CPMBR):

CPMBR = (BSJ-spanning reads supporting the candidate / total BSJ-spanning reads in the corresponding library) × $10^6$.

Cross-replicate clustering was performed by processing candidate BSJs in descending order of CPMBR. Each unassigned BSJ served as a cluster seed, and all remaining unassigned BSJs with both start and end coordinates within ± 5 nt of the corresponding seed coordinates were assigned to that cluster. Seed coordinates were retained as the representative cluster coordinates.

*Data-driven selection of abundant vcircRNAs*

Because BSJ-supporting reads represent a small fraction of viral sequencing reads and are therefore particularly sensitive to stochastic sampling, we sought to identify an abundance threshold that preferentially retained reproducible BSJ clusters while reducing replicate-specific low-support events.

The distribution of CPMBR values across the three biological replicates was at 100 thresholds corresponding to successive quantiles from 0 to 99. At each threshold, BSJ clusters were classified according to the number of biological replicates with CPMBR values at or above the threshold: one ($n_1$), two ($n_2$), or three ($n_3$). The resulting distributions are shown in Fig. S1.

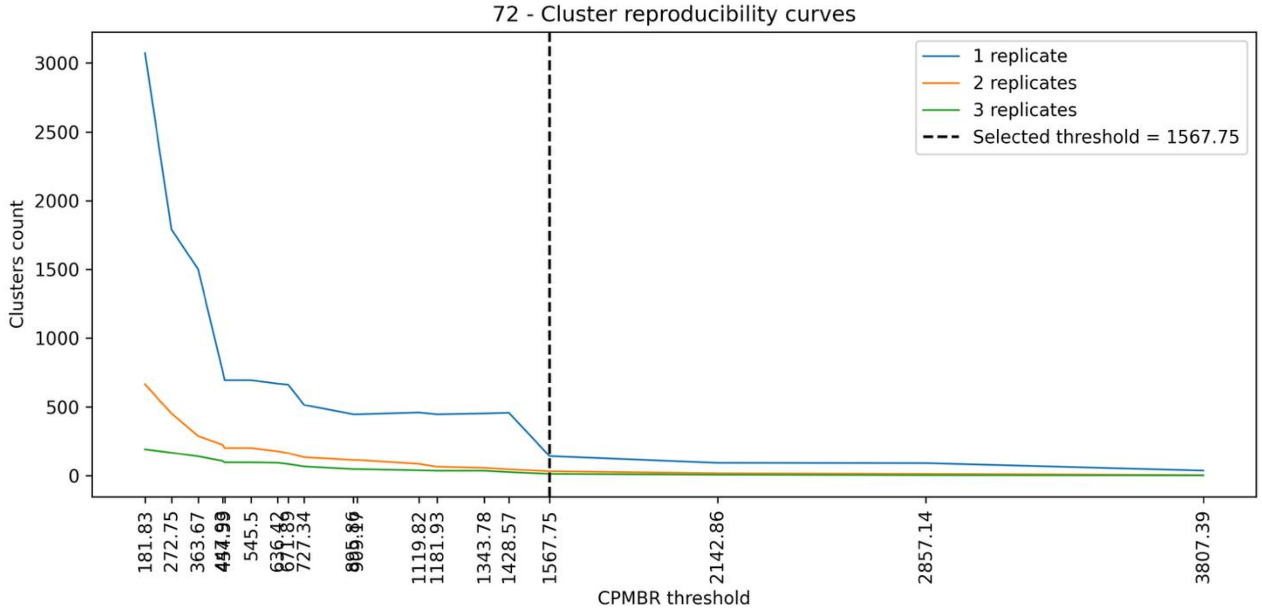


**Figure S1 - BSJ cluster reproducibility across CPMBR thresholds at 72 h post-infection.** Curves show the number of BSJ clusters with CPMBR values at or above each threshold in exactly one (blue, $n_1$), two (orange, $n_2$), or three (green, $n_3$) biological replicates. The vertical dashed line indicates the selected threshold (CPMBR = 1567.75), which maximized the step score, loss($n_1$)/[loss($n_2$ + $n_3$) + 1], among thresholds retaining at least one cluster above the threshold in two or more replicates.

For consecutive thresholds, the incremental loss in each class was calculated. To identify the threshold providing the most favorable reduction of replicate-specific events relative to reproducible events, a step score was defined as:

$$\text{step\_score} = \text{loss}(n_1) / (\text{loss}(n_2 + n_3) + 1)$$

Higher step scores therefore identify threshold increments that remove a comparatively large number of replicate-specific clusters while preserving clusters supported across multiple biological replicates. To avoid excessively stringent filtering, candidate thresholds retaining no clusters supported by at least two replicates were excluded. The threshold maximizing the step score among the remaining candidates was selected (CPMBR = 1567.75).

*Definition of the high-confidence vcircRNA core*

The selected abundance threshold was subsequently used to prioritize highly supported BSJ clusters while accounting for variability in sequencing depth, circRNA enrichment and stochastic sampling among biological replicates.

A cluster was retained if its CPMBR value reached or exceeded the selected threshold in at least one biological replicate. For each retained cluster, any non-zero BSJ support in an additional replicate contributed to the cross-replicate intersection, irrespective of whether its CPMBR value reached the selected threshold.

Retained clusters belonging to the intersection of all three biological replicates were designated core replicate-consistent vcircRNAs (core rc-vcircRNAs). These clusters were ranked by decreasing mean CPMBR across the three replicates and used for subsequent characterization and selection for experimental validation.

**RT-qPCR analysis of ZIKV replication kinetics and selected ZIKV vcircRNAs**

*RNA reverse transcription (RT)*

RT was performed on extracted RNA from NSCs and MRC5 infected cells, using SuperScriptTM III Reverse Transcriptase Kit (Thermo Fisher Scientific). The reaction was prepared in a total volume of 20 µL and it included: 1µg of RNA, 25 µM of deoxynucleotide triphosphates (dNTPs, Applied Biosystems), 50 µM of Random Hexamers (Thermo Fisher Scientific), 5X First-strand buffer, 100 mM of DTT, 20 U/µL of RNase Inhibitor (Applied Biosystems), 200 U/µL of Superscript III RT and nuclease-free water. The reaction cycle was composed of 5 minutes at 25°C, followed by 60 min at 48°C and 15 min at 70°C to inactivate the enzyme.

*Real-time PCR (qPCR)*

qPCR was performed on cDNAs from RT reaction using PowerUp™ SYBR™ Green Master Mix (Applied Biosystems). The mix was composed by 12,5 µL of master mix, 0,25 µL of Forward primer 10 µM, 0,25 µL of Reverse primer 10 µM, and 10 µL of nuclease free water. The mix was aliquoted in a 96-well plate for standard qPCR and 2 µL of RT were added to each mix. Primers used to perform qPCR experiment are reported in Table S1.

**Table S1. PCR primers for the amplification and validation of selected BSJs.**

| ***BSJ coordinates*** | ***Forward primer sequence*** | ***Reverse primer sequence*** |
|---|---|---|
| *circ_9971_10391* | 5' – ACCTATCCACCCAAGTTCGC – 3' | 5' – CCCAGTTGGAACCCAGTCAA – 3' |
| *circ_8318_8611* | 5' – CGCGGAAACGTGGTTCTTTG – 3' | 5' – GCTCGTGGTGGACACACTTT – 3' |
| *circ_8630_8909* | 5'- CCACGAGTCTGCACCAAAGA – 3' | 5' – GACTCCAGTCACCACATCCC – 3' |
| *circ_1586_1872* | 5' – GGAGGCTGAGATGGATGGTG – 3' | 5' – AGCGTGCCAAGGTAATGGAA – 3' |

| | | |
|---|---|---|
| *circ_4964_5452* | 5' – GCCACCTTCACTTCACGTCT – 3' | 5' – ATCCTGAAGTTCCTGCTGGG – 3' |
| *circ_10012_10390* | 5' – ACCTATCCACCCAAGTTCGC – 3' | 5' – TCAGTGGTCATCCATTCTCCC – 3' |
| *circ_5461_5868* | 5' – AGAGTGGGACTTTGTCGTGA – 3' | 5' – CGCCCATCTCAACCCTTGTT – 3' |
| *circ_937_1243* | 5' – GCTGCCCAACACAAGGTGAA – 3' | 5' – TGCACCTGATGCTGTATGCC – 3' |
| *circ_5039_5433* | 5' – ACAGCAGTCAATGTCACCCA – 3' | 5' – CCGTTTTTGATCACGACCCC – 3' |
| *circ_7063_7397* | 5' – GCTGGCATCATGAAGAACCC – 3' | 5' – GGCCATCGCCATTAAGGAGT – 3' |
| *circ_6859_7220* | 5' – TTGTTTGGCATGGGCAAAGG – 3' | 5' – ATTGGCGGTAATCAAGCCCA – 3' |
| *circ_7399_7723* | 5' – TGGCTTGGTCAAGAGACGTG – 3' | 5' – GACAGTATGGCGCTGGAGAC – 3' |
| *GAPDH* | 5' – GAAGGTGAAGGTCGGAGTC – 3' | 5' – GAAGATGGTGATGGGATTTC – 3' |
| E (ZIKV) | 5' – TGGGTTGATGTTGTCTTGGA – 3' | 5' – ATCTTACCTCCGCCATGTTG – 3' |

**Rolling-circle RT and Nanopore validation of candidate vcircRNAs**

*RNA enrichment and rolling-circle reverse transcription*

Prior to rolling-circle reverse transcription (RRT), total RNA was polyadenylated to facilitate RNase R-mediated depletion of linear ZIKV RNAs, which lack a 3′ poly(A) tail. Isolated RNA (15µg) from mock and ZIKV-infected samples were treated with *Escherichia Coli* Poly(A) polymerase (New England Biolabs): the reaction mix was prepared using total RNA from each condition (mock and infected), 10X Poly(A) Polymerase Reaction Buffer, 3 µL of 10mM Adenosine-5'-Triphosphate (ATP, NEB), 2 µL of E. coli Poly(A) Polymerase, 10 U RNase Inhibitor (Applied Biosystems), and nuclease-free water. The reaction was incubated at 37°C for 20 minutes. Subsequently, the RNA was purified with the RNA clean and concentrator-5 kit (ZYMO Research). Briefly, a binding buffer and ethanol were added to samples, then placed in the spin columns; two wash steps were performed and then the RNA was eluted in DNase/RNase-free water. Purified RNA was subsequently treated with RNase R (LGC Biosearch Technologies) at 4 U/µg RNA in the supplied 10× reaction buffer for 30 min at 37°C, followed by enzyme inactivation at 60°C for 10 min and purified using the RNA Clean & Concentrator Kit, as previously described.

RRT was carried out separately for each vcircRNA using candidate-specific primers located downstream of the predicted BSJ (Table S2). In each reaction, 1 µg of RNA was combined with 1 µL of 10 µM BSJ-spanning primer and 1 µL of 25 mM dNTPs, incubated at 65°C for 5 min and immediately transferred to ice for 1 min. First-Strand Buffer, 1 µL of 100 mM DTT, 20 U RNase inhibitor and 200 U SuperScript III reverse transcriptase were subsequently added. Reverse transcription was performed for 1 h at 50°C or 55°C, depending on the primer, followed by enzyme inactivation at 75°C for 15 min.

*Amplification of concatemeric RRT products and dsDNA generation*

Because RRT generates concatemeric cDNAs containing tandem copies of the vcircRNA sequence, the corresponding L-PCR primer-binding site is repeated along the resulting cDNA. Primer annealing at different copies of this repeated binding site can therefore generate amplification products encompassing one or multiple consecutive copies of the vcircRNA sequence. Single-primer linear PCR (L-PCR) was performed using Platinum Taq DNA Polymerase (Thermo Fisher Scientific). Each 60-µL reaction contained 12 µL reaction buffer, 6 µL of 2 mM dNTP mix, 1.2 µL of 10 µM candidate-specific primer (Table S1, Forward primers), 2.5 U Platinum Taq DNA Polymerase and 6 µL of the RRT reaction. Thermal cycling consisted of an initial denaturation at

94°C for 10 min, followed by 40 cycles of 94°C for 30 s, 60°C for 30 s and 68°C for 1 min, with a final extension at 68°C for 7 min.

Following single-primer amplification, a second amplification step was performed using a candidate-specific reverse primer (Table S1, Reverse primers) to generate double-stranded DNA (dsDNA) molecules suitable for Nanopore library preparation. This step was intended to convert the products generated during the first L-PCR into dsDNA while preserving their variable concatemeric lengths and, consequently, the number of consecutive vcircRNA copies represented within each molecule.

Before this step, all L-PCR were pooled together and purified using the DNA Clean & Concentrator Kit (Zymo Research). Second amplification reactions were performed using a reaction mixture containing 0,25 µL of each BSJ-spanning primer (10 µM), 0,4 µL of Platinum Taq Hot start (5 U/µL), 12 µL of 5X Platinum II PCR Buffer, 5 µL of 2 µM dNTPs and 10,4 µL of nuclease free water. The single PCR step consisted of 30 seconds at 94 °C, 30 seconds at 60 °C and 2 minutes at 68 °C. The resulting dsDNA products were purified using the DNA Clean & Concentrator Kit. Purified products were subsequently treated with Klenow enzyme by addition of 10× NEB2 buffer, Klenow enzyme at 1 U/µg DNA, BSA at a final concentration of 0.2 mg/mL and 40 µM dNTPs in a final volume of 50 µL. Reactions were incubated at 37°C for 15 min, followed by enzyme inactivation at 65°C for 10 min and a final purification using the DNA Clean & Concentrator Kit.

**Table S2. PCR primers used for BSJ-spanning rolling circle reverse transcription (RRT).**

| ***BSJ coordinates*** | ***RRT primer sequence*** |
|---|---|
| *circ_9971_10391* | 5' – TCTGTGGAAATAAGATTGGTGCT – 3' |
| *circ_8318_8611* | 5' – GTACATCTCATGTGTGGGGGCCT – 3' |
| *circ_8630_8909* | 5' – AACCCCGTTTATTAAATATTGCC – 3' |
| *circ_1586_1872* | 5' – TCATAGTCAAGTAAGTTTATCCATT – 3' |
| *circ_4964_5452* | 5' – CCATCCTTTGTCTTATAGTTGGGG – 3' |
| *circ_10012_10390* | 5' – CACAGATGAACAAGATTGGTGCT – 3' |
| *circ_5461_5868* | 5' – CTCATCCATAATAAAGTTGGCG – 3' |
| *circ_937_1243* | 5' – ACCAAGTATATGAGTGTCTGATTG – 3' |
| *circ_5039_5433* | 5' – CTCCCACACTTGGCTGTA – 3' |
| *circ_7063_7397* | 5' – CTGGGGTAATGTCATTGT– 3' |
| *circ_6859_7220* | 5' – GCCATTTGGTTGAGTAGCA – 3' |
| *circ_7399_7723* | 5' – TGGGGGTCAAATCTGGTTC – 3' |

*Nanopore library preparation and sequencing*

Pooled dsDNA products were prepared for Nanopore sequencing using the Native Barcoding Kit 24 V14 (SQK-NBD114.24; Oxford Nanopore Technologies) according to the manufacturer's instructions. Short Fragment Buffer (SFB) was used during the post-adapter-ligation clean-up step to retain DNA fragments across the complete size range. The resulting library was loaded onto an R10.4.1 flow cell (FLO-MIN114) and sequenced for approximately 18 h using a MinION Mk1D device.

**Bioinformatic analysis of Nanopore validation reads**

*Construction of candidate-specific concatemeric references*

Nanopore reads generated following rolling-circle reverse transcription and linear PCR amplification were analyzed using candidate-specific custom reference sequences designed to represent the expected structure of rolling-circle products.

For each candidate vcircRNA, the predicted circular sequence was reconstructed from its genomic coordinates and concatenated in tandem ten times. The corresponding linear-PCR primer sequence was additionally appended to the 5′ end of the concatemeric reference to reproduce the expected starting structure of the amplified Nanopore reads.

The use of tandem references allowed individual long reads to align continuously across multiple consecutive copies of the predicted circular sequence and therefore enabled direct quantification of repeated BSJ-spanning events within individual molecules.

*Nanopore read alignment and filtering*

Nanopore reads were aligned against the candidate-specific concatemeric references using minimap2 v2.30 with the lr:hq preset. Resulting SAM alignments were partitioned according to reference sequence so that reads corresponding to each vcircRNA candidate could be evaluated independently.

Only mapped primary alignments were retained for downstream analysis. Reads had to meet a candidate-specific minimum expected length, determined by the linear-PCR primer position and the predicted vcircRNA length. Retained alignments were required to cover at least 90% of the query sequence and have at least 90% sequence identity, defined as the fraction of exact matches across all alignment positions, including mismatches, insertions, and deletions.

*Quantification of repeated BSJ-spanning events*

Because both the custom references and the rolling-circle RT-derived reads contain tandem repeats of the same predicted vcircRNA sequence, alignments can be shifted by one or more repeat units without changing their sequence correspondence. Absolute alignment coordinates are therefore not uniquely defined, and BSJ support was evaluated in a normalized coordinate system defined relative to the reverse-transcription primer binding site.

For each retained read, the aligned sequence was evaluated to determine the number of complete traversals of the predicted circular RNA sequence. A complete traversal was counted when the read extended through the expected BSJ and continued for at least 5 nt beyond the junction. The 5-nt extension requirement was introduced to ensure that BSJ support reflected sequence information extending beyond the junction rather than termination immediately at the predicted breakpoint.

The number of complete BSJ-spanning events was determined independently for each read. Reads containing multiple consecutive BSJ-spanning events were interpreted as concatemeric products generated by rolling-circle reverse transcription and therefore provided strong evidence supporting the circular nature of the corresponding RNA molecule.

### 1. Results

**Table S3. Deduplicated read-pair alignment statistics for ZIKV-infected neural stem cell samples collected at 24, 48, and 72 hours post-infection (hpi).** For each sample, the total number of read pairs and the numbers and percentages aligning to the human T2T reference and the ZIKV reference genome (KX377337.1) are reported.

| SAMPLE | | | ALIGNMENT STATS (DEDUP) | | |
|---|---|---|---|---|---|
| REP | HPI | ID | TOTAL | HOST (T2T) | ZIKV (KX377337.1) |

| | | | read pairs | read pairs | %TOTAL | read pairs | %TOTAL |
|---|---|---|---|---|---|---|---|
| **A** | **24** | **A3** | 41,598,692 | 34,033,768 | 81.81% | 39,580 | 0.10% |
| **B** | **24** | **B3** | 23,344,769 | 23,254,957 | 99.62% | 47,691 | 0.20% |
| **C** | **24** | **C3** | 25,059,925 | 24,966,485 | 99.63% | 62,732 | 0.25% |
| **A** | **48** | **A6** | 77,197,246 | 72,997,826 | 94.56% | 165,169 | 0.21% |
| **B** | **48** | **B6** | 26,520,332 | 26,304,462 | 99.19% | 187,711 | 0.71% |
| **C** | **48** | **C6** | 24,831,146 | 24,534,736 | 98.81% | 248,608 | 1.00% |
| **A** | **72** | **A9** | 75,001,435 | 69,580,324 | 92.77% | 929,342 | 1.24% |
| **B** | **72** | **B9** | 25,491,584 | 25,025,009 | 98.17% | 411,335 | 1.61% |
| **C** | **72** | **C9** | 23,679,575 | 22,735,780 | 96.01% | 872,714 | 3.69% |

**Table S4. Back-spliced read-pair counts and strand distribution in ZIKV-infected neural stem cell samples collected at 24, 48, and 72 hours post-infection (hpi).** A read pair was counted as back-spliced if at least one mate spanned a back-splice junction (BSJ). The total percentage is calculated relative to the ZIKV-aligned read pairs in each sample, reported in Table S3; the positive- and negative-strand percentages are calculated relative to all back-spliced read pairs in that sample. The final column reports the number of unique BSJs supported by at least one back-spliced read pair.

| **SAMPLE** | | | **CUSTOM BSJ-DETECTION WORKFLOW** | | | | | | |
|---|---|---|---|---|---|---|---|---|---|
| | | | **BACK-SPLICED READS** (at least one mate is a BSJ-spanning read) | | | | | | **circRNAs** (number of unique junctions supported at least by 1 count) |
| | | | **TOTAL** | | **POSITIVE STRAND (+)** | | **NEGATIVE STRAND (-)** | | |
| **REP** | **HPI** | **ID** | *read pairs* | *%ZIKV* | *read pairs* | *% BSJ-SPANNING READS* | *read pairs* | *% BSJ-SPANNING READS* | *absolute number* |
| **A** | **24** | **A3** | 495 | 1.25% | 446 | 90.10% | 49 | 9.90% | 481 |
| **B** | **24** | **B3** | 356 | 0.75% | 317 | 89.04% | 39 | 10.96% | 350 |
| **C** | **24** | **C3** | 645 | 1.03% | 590 | 91.47% | 55 | 8.53% | 633 |
| **A** | **48** | **A6** | 3,997 | 2.42% | 3,649 | 91.29% | 348 | 8.71% | 3,813 |
| **B** | **48** | **B6** | 2,071 | 1.10% | 1,785 | 86.19% | 286 | 13.81% | 1,975 |
| **C** | **48** | **C6** | 6,152 | 2.47% | 5,605 | 91.11% | 547 | 8.89% | 5,749 |
| **A** | **72** | **A9** | 20,705 | 2.23% | 18,959 | 91.57% | 1,746 | 8.43% | 17,984 |
| **B** | **72** | **B9** | 8,264 | 2.01% | 7,588 | 91.82% | 676 | 8.18% | 7,428 |
| **C** | **72** | **C9** | 38,722 | 4.44% | 35,718 | 92.24% | 3,004 | 7.76% | 31,532 |

**Figure S2**

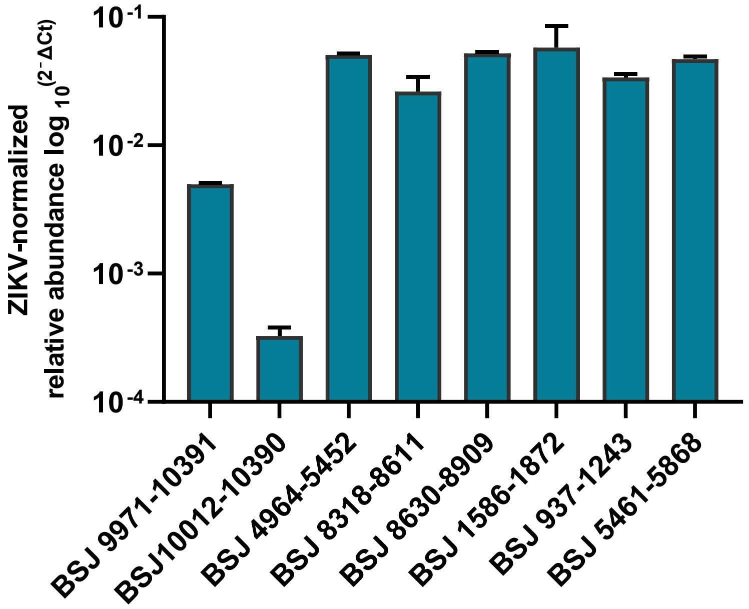


**Figure S2. Candidate core replicate-consistent vcircRNAs (rc-vcircRNAs) validation in MRC-5 cells by quantitative reverse-transcription PCR (RT-qPCR).** The bar graph shows the relative abundance of candidate core rc-vcircRNAs, expressed as $\log_{10}{}^{(2^{-\Delta Ct})}$, with Ct values normalized to ZIKV genomic RNA, quantified using primers targeting the E gene. Data show the mean ± standard deviation (SD) of n = 3 independent experiments.

**Figure S3**

| vcircRNA ID | Predicted length (nt) | Genomic region | rc-vcircRNAs expression rank position | Junction sequence | Validations support (consecutive BSJ-spanning read count) x1 | x2 | x3 | x4 | x5 |
|---|---|---|---|---|---|---|---|---|---|
| ZIKV:5039-5433 | 395 | NS3 | n.a. | AGCCAA \| GTGTGG | | | | | |
| ZIKV:7063-7397 | 335 | NS4B | n.a. | ATGACA \| TTACCC | | | | | |
| ZIKV:6859-7220 | 362 | NS4B | n.a. | TACTCA \| ACCAAA | | | | | |
| ZIKV:7399-7723 | 325 | NS4B | n.a. | CCAGAT \| TTGACC | | | | | |

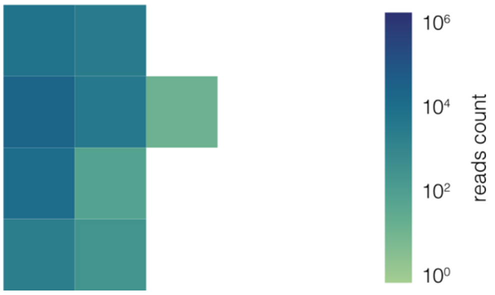


**Figure S3. Summary of four additional vcircRNAs validated by rolling-circle RT and Nanopore sequencing.** These candidates were excluded from the core set of rc-vcircRNAs because they did not meet the stringent inclusion criteria. The table reports their genomic coordinates, predicted lengths, genomic regions, and back-splice junction (BSJ) sequences. The heatmap shows read counts by the number of consecutive BSJ crossings (one to five), with counts represented on a logarithmic color scale.
